# *In silico* characterization of cysteine-stabilized αβ defensins from neglected unicellular microeukaryotes

**DOI:** 10.1101/2022.10.13.512120

**Authors:** Marcus Vinicius Xavier Senra

## Abstract

**Background:** The emergence of multi-resistant pathogens have increased dramatically in recent years, becoming a major public-health concern. Among other promising antimicrobial molecules with potential to assist in this worldwide struggle, cysteine-stabilized αβ (CS-αβ) defensins are attracting attention due their efficacy, stability, and broad spectrum against viruses, bacteria, fungi, and protists, including many known human pathogens.

**Results:** Here, 23 genomes of ciliated protists were screened and three CS-αβ defensins with a likely antifungal activity were identified and characterized using bioinformatics from two freshwater and culturable species *Laurentiella* sp. (LsAMP-1 and LsAMP-2) and *Euplotes focardii* (EfAMP-1). Although any potential cellular ligand could be predicted for LsAMP-2 and EfAMP-1; evidences from structural, molecular dynamics, and docking analyses suggest that LsAMP-1 may form stably associations with phosphatidylinositol 4,5-bisphosphates (PIP2), a phospholipid found on many eukaryotic cells, which could, in turn, represent an anchorage mechanism within plasma membrane of targeted cells.

**Conclusion:** These data stress that more biotechnology-oriented studies should be conducted on neglected protists, such ciliates, which could become valuable sources of novel bioactive molecules for therapeutic uses.

## Background

The discovery and therapeutic use of antimicrobial agents represent one of the greatest milestones in medical science, contributing to saving countless lives every year [1]. Unfortunately, only 7 years were required, since penicillin introduction [2], for the emergence of an antibiotic-induced resistant strain (*Staphylococcus aureus*), which were isolated from hospitalized patients in London in 1948 [3]. Afterwards, countless preoccupying cases of resistant and multidrug-resistant microorganisms, such as *Mycobacterium tuberculosis [4]*, *Streptococcus pyogenes [5]*, *Staphylococcus aureus [6]*, *Enterobacter cloacae [7]*, *Pseudomonas aeruginosa [8]*, *Candida albicans*, and *Aspergillus fumigatus [9]* started to be reported, resulting from natural selection processes, but exacerbated from the misuse and overuse of antibiotics [2]. Therefore, the development of new antimicrobials is utmost necessary to combat this major concern in public health.

One promising class of molecules is the cysteine-stabilized α-helical β-sheet (CS-αβ) defensins. These antimicrobial peptides (AMPs) are typically small (34-54 amino acid residues), amphipathic (defined hydrophobic and hydrophilic regions within the same molecule), mostly cationic, and cysteine-rich molecules that adopt a canonical CS-αβ scaffold, consisting in an α-helix and two β-strand antiparallel β-sheet stabilized by two disulfide bridges linking the α-helix to the C-terminal β-strand, and a third bridge connecting the N-terminus to the first β-strand [10].

Critical players of humoral defense systems of fungi [11], plants [12] and some invertebrates [13, 14], these AMPs may exert their functions through a diversity of mechanisms of actions, including membrane disruption and pore formation [15], ion channels blockage [16, 17], and interference to different intracellular pathways [18–20]; and are highly effective against a broad spectrum of pathogens [21]. While plant CS-αβ defensins are predominantly antifungal [21], with a few insecticidal [18] and bactericidal [21]; fungal and animal CS-αβ defensins are mostly antibacterial [21], with some reports of antiprotozoals [22, 23] and fungi [13, 24, 25].

Here, 23 publicly available genomes of ciliated protists, an ubiquitous and highly diverse group of unicellular microeukaryotes [26], were screened for CS-αβ defensin homologs and possible cellular ligands are presented.

## Results

Screenings for CS-αβ defensin homologs within the predicted proteomes of 23 ciliate species were performed using a combined approach, which consisted in pairwise sequence alignments against SwissProt database and profile search, using a previously described CS-αβ defensins cysteine distribution pattern that is conserved across CS-αβ defensins [27]. After a series of filtering steps (see methods for details), three unannotated CS-αβ defensins were identified from two freshwater hypotrichous ciliates, *Laurentiella* sp. (LsAMP-1 and LsAMP-2) and *Euplotes focardii* (EfAMP-1) and data are summarized in Table 1.

**Table 1.**
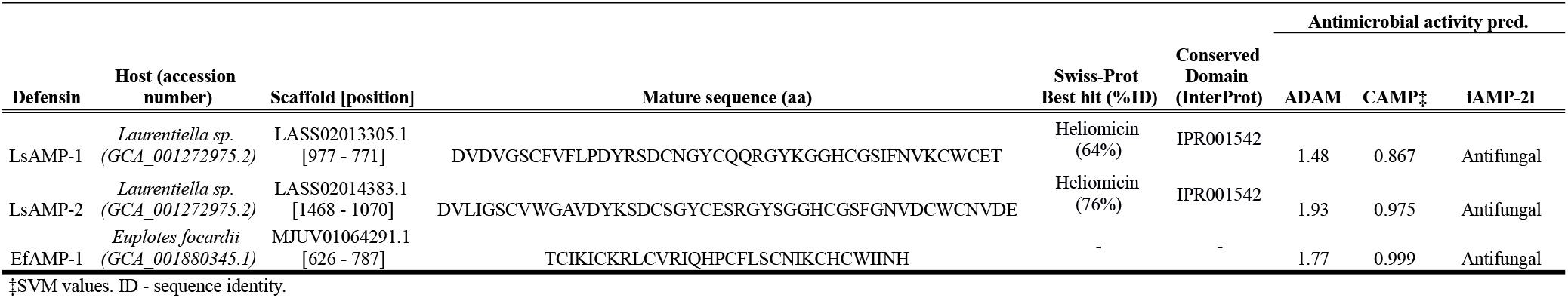
Basic sequence information and antimicrobial activity prediction of ciliate defensins.

LsAMP-1 and LsAMP-2 are both anionic (net charge of −1 and −5 at pH 7,0) with N-terminal signal peptides spanning through the first 20 amino acid (aa) residues and mature sequences of 44 and 46 aa residues, in which contains a conserved invertebrate defensin domain (IPR001542) (Table 1). These defensins share a 61% of sequence identity and are highly similar to previously characterized lepidopterans antifungal CS-αβ defensins, heliomicin (PDB:1i2u/1i2v), isolated from *Heliothis virescens* (Noctuidae) [24] (64% and 76% of identity); and ARD1 (PDB:1p0a/1p0o), isolated from *Archaeoprepona demophon* (Nymphalidae) [28] (59% and 71% identity) (Table 1 and Fig. 1).

**Fig. 1.**
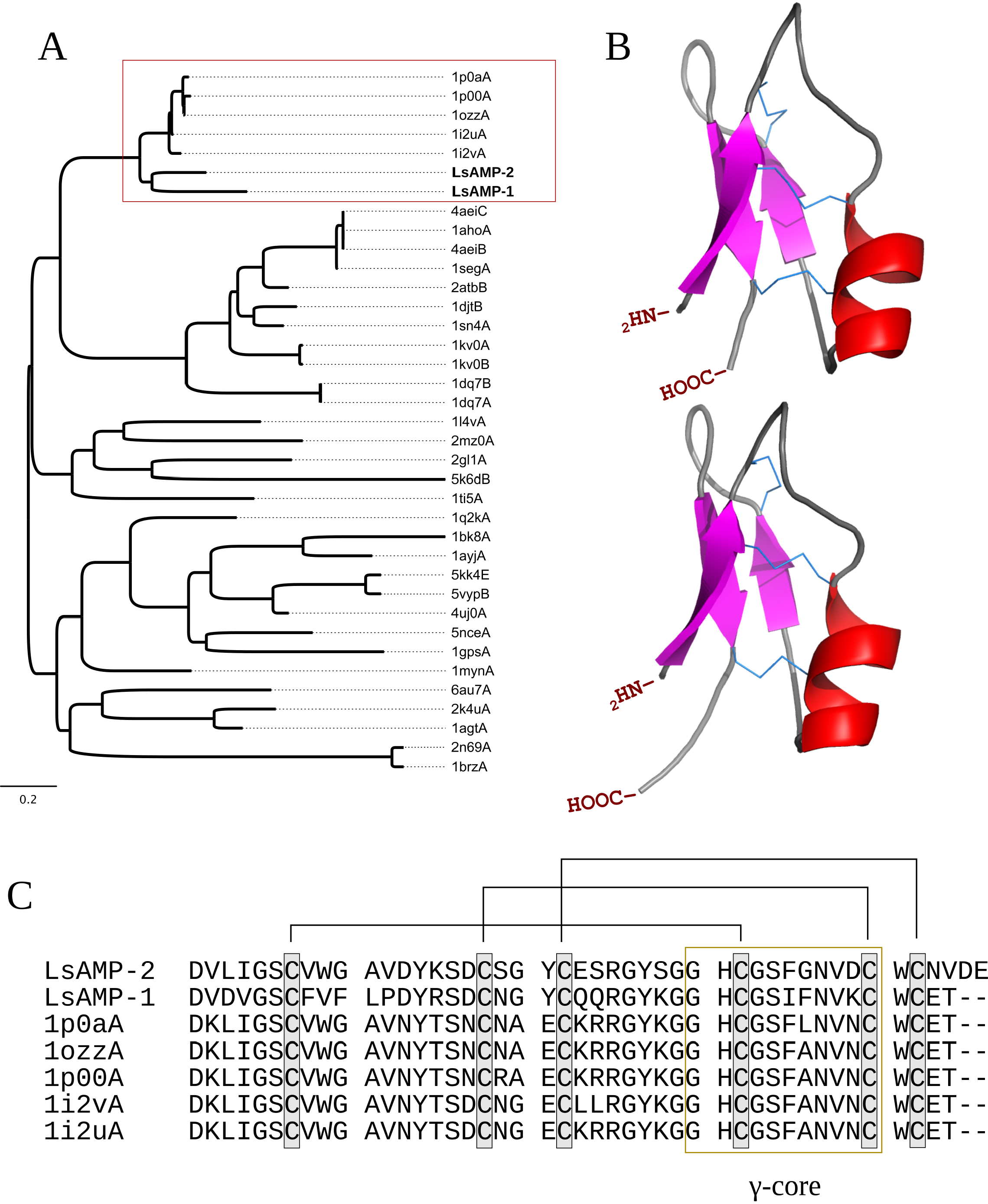
(A) Neighbor-joining phylogenetic tree inferred from structural alignment of proteins available from PDB. LsAMP-1 and LsAMP-2 are in bold. The red square highlights the monophyletic cluster including LsAMP-1, LsAMP-2, heliomicin (1i2vA and 1i2uA) and ARD1 (1p0aA and 1p0oA). (B) Theoretical three-dimensional models of LsAMP-1 (top) and LsAMP-2 (botton). Disulfide bridges are connected with sticks. And (C) alignment of defensin mature sequences highlighted in (A). Lines in black represent the cysteine connections and the γ-core of these proteins are within the yellow square

LsAMP-1 and LsAMP-2 theoretical structures were predicted through comparative modeling, using heliomicin and ARD1 as templates, respectively (Fig. 1b and Table 2), both consisting of an N-terminal β-strand followed by an α-helix and two antiparallel β-strands (βαββ) scaffold, stabilized by three disulfide bridges (Cys^1^-Cys^4^, Cys^2^-Cys^5^, and Cys^3^-Cys^6^) (Fig. 1b-c). At their C-terminus there is a canonical signature (Gly-X-Cys-X_3-9_-Cys, where X represents any residue) recognized as the γ-core, which plays a role in the antimicrobial activity of other CS-αβ defensins [29, 30] (Fig. 1c).

**Table 2.** Template information and quality check of ciliate defensin 3D models.

| Defensin | Template | Template classification | ID† | Cov‡ | Ramachandran plot analysis |  | Outliers | ProSA (Z-Score) |
| --- | --- | --- | --- | --- | --- | --- | --- | --- |
|  |  |  |  |  | Residues in favored regions (%) | Residues in allowed regions (%) |  |  |
| LsAMP-1 | 1i2uA | Antifungal | 0,64 | 1 | 92,9 | 100 | - | -5,43 |
| LsAMP-2 | 1p0aA | Antifungal | 0,71 | 0,97 | 90,9 | 100 | - | -5,74 |
| EfAMP-1 | 1ayjA | Antifungal | 0,28 | 0,97 | 93,3 | 100 | - | -5,3 |

† Sequence identity (ID) and ‡ coverage (cov) ‡ between query/template.

**Table 3.** Predicted cellular ligand of LsAMP1.

| Model | Predicted ligand | PDB hit | Cofactor |  |  | DockThor |  |
| --- | --- | --- | --- | --- | --- | --- | --- |
|  |  |  | BS-score* | Identity | Coverage | Binding energy (-Kcal/mol) | Hydrogen bonds |
| LsAMP-1 | PIP2 | 4cqkA | 1.10 | 0.21 | 0.82 | 7.950 | Lys-39,Asn-3 |

\*BS-score is a measure provided of local similarity between template and predicted binding site in the defensins.

On the other hand, EfAMP-1 is a cationic CS-αβ defensin (net charge of +5 at pH 7,0) with an N-terminal signal peptide spanning the first 22 aa residues and a mature peptide of 32 aa residues (Table 1). Differently to what was found for LsAMP-1 and LsAMP-2, EfAMP1 seems rather divergent, with no significant sequence identity (>30%) to proteins in PDB and no obvious InterProt conserved domains (Table 1). In this case, fold recognition analysis had to be used to identify a plant antifungal CS-αβ, RS-AFP1[31], isolated from the Magnoliophyta *Raphanus sativus* (Brassicales), which shares a significant structural alignment (c-score >1) with EfAMP-1, as a template to model the 3D structure of this defensin (Table 2). This spatial similarity was further supported through phylogenetic analysis, where EfAMP-1 emerges in a cluster with different plant CS-αβ defensins (Fig. 2a).

**Fig. 2.**
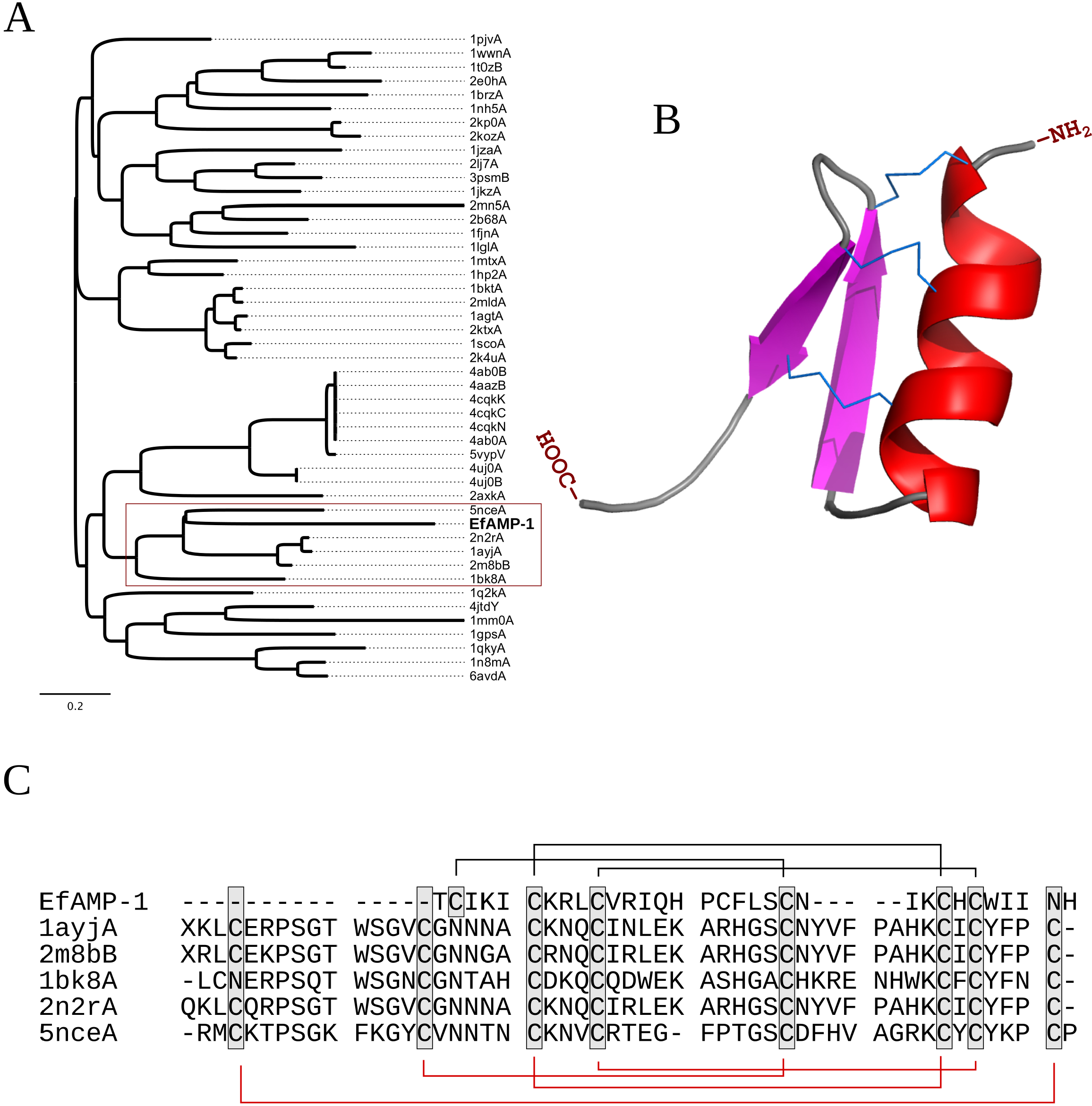
(A) Neighbor-joining phylogenetic tree inferred from structural alignment of proteins available from PDB. EfAMP-1 is in bold. The red square delimits a monophyletic cluster of plant CS-αβ defensins related to EfAMP-1. (B) Theoretical three-dimensional model of EfAMP-1. Disulfide bridges are connected with sticks. And (C) alignment of defensin mature sequences highlighted in (A). Lines in black represent the cysteine connections predicted for EfAMP-1 and lines in red the connections described for the plants defensins

EfAMP-1 theoretical structure (Fig. 2b) shows an N-terminal α-helix and two antiparallel β-strand (αββ) scaffold, stabilized by three cysteine bridges (Cys^1^-Cys^5^, Cys^2^-Cys^6^, and Cys^3^-Cys^7^), consisting of two connections between the α-helix and the second β-strand and one linkage between the N-terminal to the first β-strand (Fig. 2c), which is typically seen in CS-αβ motifs [27].

All amino acid residues from these three defensin models emerged within favorable and acceptable regions of Ramachandran plots (Table 2); and analyzes of root mean square deviations (RMSD), root mean square fluctuation (RMSF), and dictionary of protein secondary structure (DSSP) indicate that most movements in amino acid residues are within acceptable ranges (Fig. 3) and secondary structures remained almost unchanged along the entire MD simulation (Supplentary material 1), collectively evidencing the quality and stability of these models.

**Fig. 3.**
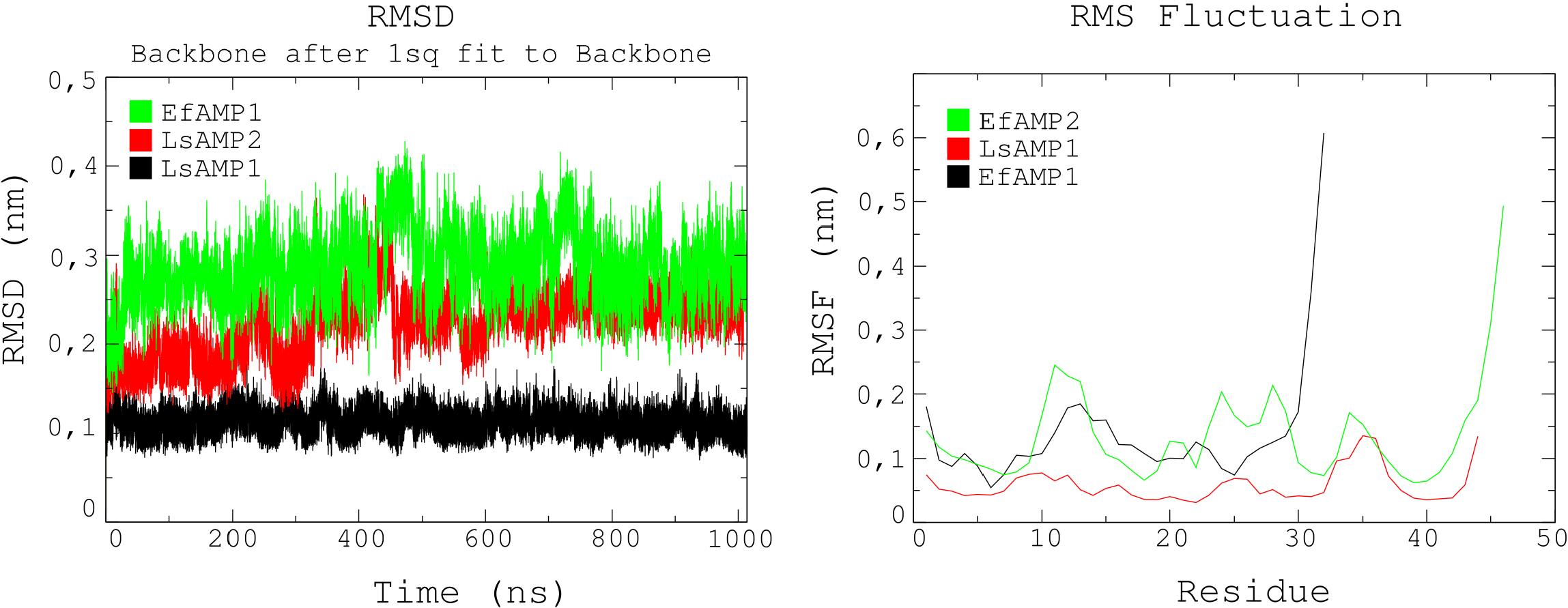
Molecular dynamic evaluation of LsAMP-1, LsAMP-2, and EfAMP-1 theoretical structures. A root mean square deviation (RMSD) plot, showing the mean backbone variation over the 1μs of this run

Once in hands with high-quality models of these defensins, the next move was to evaluate possible cellular ligands, which was performed by combining data from structural, molecular docking, and MD simulations. Although clear candidates could not be identified for LsAMP-2 and EfAMP-1, comparative structural analysis using *Cofactor* indicates that LsAMP-1 may likely bind to phosphatidylinositol 4,5-bisphosphate (PIP2), with a BS-score of 1.10, within a surface neighboring the γ-core motif (Fig. 4). This data was further confirmed through molecular docking, where the best binding pose shows a strong energy affinity of −7.95 Kcal/mol (Fig. 4); and through a 200 ns MD simulation, which indicate that, after an short period (5 ns) of molecular accommodation, LsAMP-1 and PIP2 remained stably connected by two hydrogen bonds, linking Arg-39 and Asp-3 residues to PIP2 bisphosphate and by hydrophobic interactions between Phe-8 and Phe-10 residues and the fatty acid tail until the end of the run (Fig. 4).

**Fig. 4.**
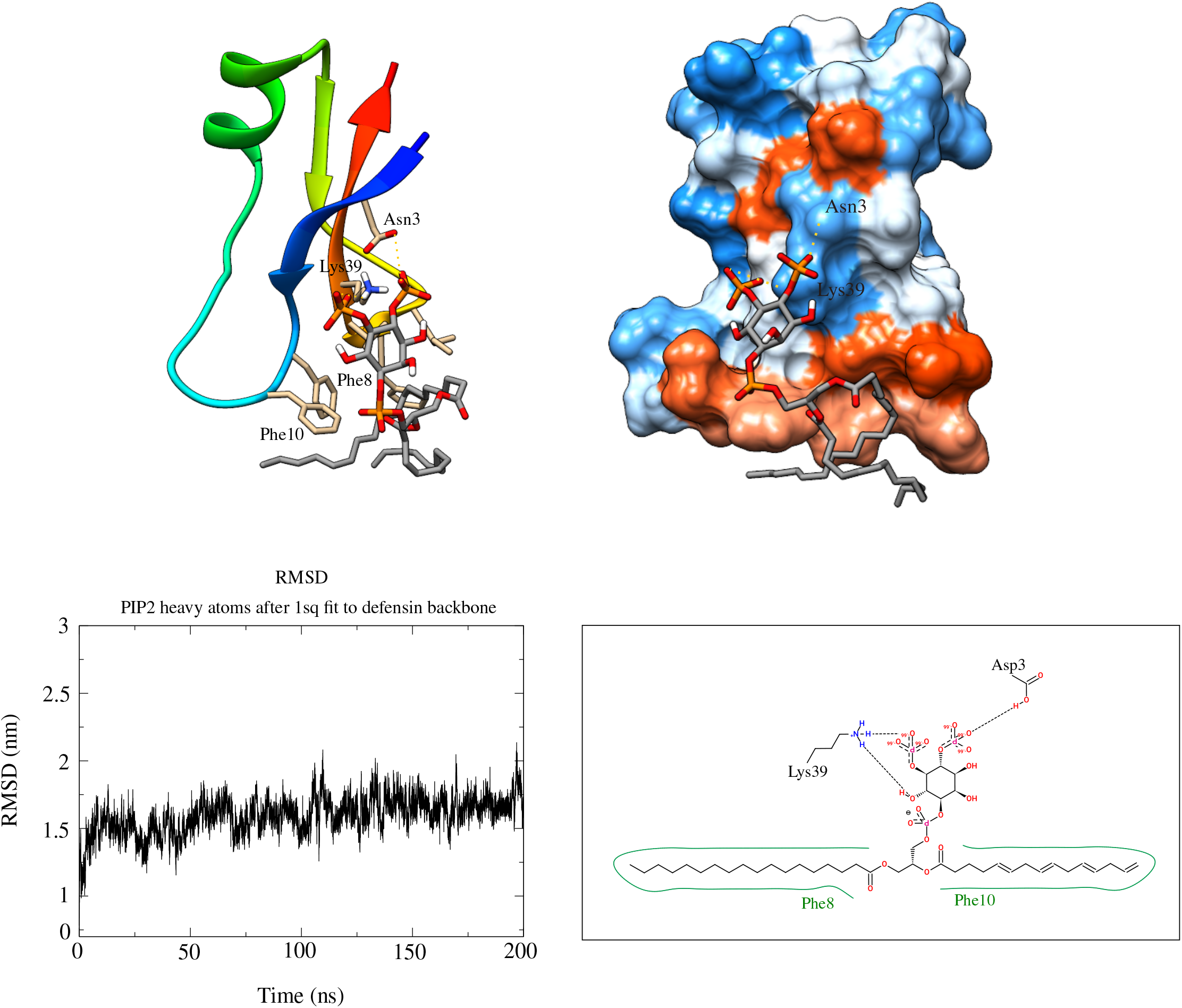
Prediction of cellular ligands of LsAMP-1. The table at the top shows data from the structural and molecular docking analyses, which suggest PIP2 as a likely ligand for LsAMP-1. At the center, 3D representations of LsAMP-1-PIP2 binding pose at the last frame of the MD simulation are presented. LsAMP-1 is depicted as new-cartoon format (left-side,) and as its hydrophobic surface (right-side), in which is colored from red to blue according to hydrophobicity, where red represents hydrophobic residues. In both cases, hydrogen bonds are marked with dashed lines. At the bottom, a root mean square deviation (RMSD) plot of the defensin mean alpha carbon variation against heavy atoms of PIP2 across 200 ns of MD simulation (left-side); and a 2D representation of LsAMP-1/PIP2, indicating hydrogen bonds (dashed lines) and hydrophobic interactions (green lines) (left-side)

## Discussion

CS-αβ defensins are ancient molecules, which may have evolved from bacterial cysteine-stabilized α-helical motifs [32] and emerged, according to the current knowledge, at the dawn of eukaryotic cell evolution, in a putative common ancestor of plants, fungi, and animals [33]. AMPs are effective against a wide variety of pathogens through different mechanisms of action [21] and exhibit low cytotoxic effects to human cells, are thermally and proteolytically stable, amenable to rational engineering, and rarely induce acquired-resistance in comparison to conventional antibiotics [21, 34]. Additionally, many CS-αβ defensins, such as lucifensin [35], isolated from the blowfly *Lucilia sericata*, and scedosporisin [36], isolated from the fungus *Scedosporidium apiospermum* are highly effective against methicillin-resistant *Staphylococcus aureus* and vancomycin-resistant Enterococci, respectively, highlighting these molecules have great potential to serve as alternatives to conventional antibiotics in human therapeutics. However, different methodological challenges still limit their practical pharmacological applications [37], such as low yields after purification procedures from natural hosts; and misfolding, degradation, and toxicity issues when total synthesis and/or heterologous expression approaches are applied [37].

Here, three novel CS-αβ defensins were characterized from two freshwater spirotrich ciliates, *Laurentiella* sp. and *Euplotes focardii*, representing the first report of such molecules in protists. This data indicates the evolutionary origin of defensins may be older than previously thought, possible within the last eukaryotic common ancestor (LECA). Moreover, many ciliates, *Laurentiella* sp. and *E. focardii* included, are culturable under *in vitro* conditions, producing dense cultures in short time spans; and are known to synthesize a variety of antibacterial, antifungal, and antiviral compounds [38–41]; I would like to stress the great and underexplored biotechnological potential these organisms as sources of novel bioactive compounds for human therapeutics.

Given the great divergence, typically observed even when comparing CS-αβ defensins from close related hosts [33], I was, in fact, expecting to find highly divergent structures within the genome of these neglected microeukaryotes, such as EfAMP-1, which is considerably shorter (N-terminal β-strand is absent) with only 3 cysteine bridges (instead of 4) in comparison to plant CS-αβ defensins within the same phylogenetic cluster (Fig. 2a-c). However, different lines of evidence suggest this may be a real CS-αβ defensin: (i) it present recognizable secretion signals and predicted antimicrobial activity (Table 1); (ii) short length (36 aa residues), cationic (net charge of +5 at pH 7.0), and amphipathic protein, with four hydrophobic amino acid residues on the same surface (I^5^, L^9^, L^19^ and I^23^), which suggests a possible region for interaction with negatively charged membranes (Fig. 2 and Table 1); (iii) presence of the canonical αββ motif (Fig. 2b-c); and also (iv) shares significant structural similarities with previously characterized plant CS-αβ defensin [31] (Table 2 and Fig. 2a).

On the other hand, LsAMP-1 and LsAMP-2 are clearly homologs of heliomicin and ARD1, two CS-αβ defensins isolated from lepidopteran insects. Because of the great phylogenetic distances between ciliates and insects, I checked for insertion elements and/or insect genes in the vicinity of these sequences. However, any sign of horizontal gene transfer events or insect DNA contamination could be noticed (data not shown), suggesting these pronounced similarities may be a consequence of strong selection pressures. Moreover, LsAMP-1 and LsAMP-2 were identified from the same host and are also greatly similar to each other (Fig. 1), raising the hypothesis that they may have evolved through duplication followed by diversification events.

Most CS-αβ defensins (including heliomicin and ARD1) characterized so far are cationic, which is often invoked to explain their selectiveness against negatively charged membranes of bacteria and fungi [42]. On the other hand, existence of many anionic CS-αβ defensins [43–46], such as LsAMP-1 and LsAMP-2 (net charge of −1 and −5 at pH 7.0, respectively), and AfusinC isolated from *Aspergillus fumigatus*, which is active against different multi-resistant humans bacterial pathogens [47], stress that electrostatic interactions may not be enough to describe the diversity of membrane-CS-αβ defensin associations.

Structural, sequence, and phylogenetic analyses suggest that LsAMP-1, LsAMP-2, and EfAMP-1 might be true defensins with fungicide activity (Table 1, Fig. 1 and 2). This data should be confirmed through future *in vitro* testes, but bring new perspectives and possibilities for the control of common resistant and multiresistant humans pathogens, such as *Aspergillus fumigatus* and *Candida auris*, which figures in the 2019 US CDC’s antibiotic resistance threats report (https://www.cdc.gov/DrugResistance/Biggest-Threats.html).

CS-αβ defensins may exert their functions through many different mechanisms of actions, which include depolarization and pore formation within targeted plasma membranes, but also could attack intracellular components, such as ones involved in protein synthesis [21]. Sequence and structural analyses performed here, indicate that LsAMP-1 may interact with PIP2, which is a rare, but essential phospholipid of eukaryotic plasma membranes, found in the inner leaflet, where it is involved in actin organization, membrane trafficking and ion channel regulations [48], and in the outer leaflet, with role in cell adhesion and motility [49]. In fact, it was previously characterized that, through interaction with PIP2, the plant defensin NAD1 forms an oligomeric arrangement in the plasma membrane of fungi and tumor cells, which is critical to induce their permeabilization [50]. I predict that there might be similarities between the mechanism of action of NAD1 and LsAMP-1, as in the best binding pose, LsAMP-1 hydrophobic region is oriented toward PIP2 long fatty acid tail, suggesting this association may facilitate the anchorage into the plasma membrane of targeted cells.

## Conclusion

This study stresses the importance of public sequence databases and computational-based methods in drug discovery, offering a rapid, efficient, and low-cost framework for the identification of new antimicrobial agents. The three CS-αβ defensins (LsAMP-1, LsAMP-2, and EfAMP-1) characterized here were identified from two fast-growing freshwater ciliates, which amenable for *in vitro* culturing, stressing the biotechnological potential of these protists, which could become a new source of biomedical molecules.

## Methods

### - *Genomic screenings for CS-αβ defensins* homologs

The 23 ciliate genomes analyzed in this study were directly retrieved from the NCBI nucleotide database [51]. Open reading frames (ORFs) were identified and extracted using *getorf* [52], applying proper genetic codes for each ciliate species, as suggested through analyzes using *facil [53]*. Since CS-αβ defensins are typically small proteins, only ORFs coding for proteins within 10-100 aa were considered in this analysis, which were performed using two combined approaches: homology search, using *BlastP [54]*, against the SwissProt database [55] applying an e-value cutoff of 10^−5^; and by pattern search, using *fuzzpro [52]* with the previously described CS-αβ defensin profile (CX2-18CX3CX2-10[GAPSIDERYW]X1CX4-17CXC, where C is for cysteine, X, stands for any amino acid residue, the subscript values are the range of occurrence and the square brackets delimit an ambiguous region in which only one of these residues can be find in that position) [27]. Next, candidates containing signal peptides - *SignalP* v5.0 [56] - (which were removed subsequently), no transmembrane domains - *Phobius* v1.01 [57] - and presenting conserved CS-αβ defensin secondary structures (αββ or βαββ) - *Psipred [58]* were selected and tested for antimicrobial activities by using 3 different AMPs predictors: *ADAM [59]*, *CAMP [60]*, and *iAMP-2l [61]*. In this framework, only sequences unanimously classified as AMPs were selected for further characterizations to reduce false-positives.

### - Protein 3D structure prediction

Templates for comparative three-dimensional (3D) modeling of LsAMP-1 and LsAMP-2 were selected from Protein Data Bank (PDB) (https://www.rcsb.org/), using *Hhpred [62]*. Then, 100 models for each protein were constructed using *Modeller* v9.19 [63], and the structure with the lowest DOPE (Discrete Optimized Protein Structure) score was selected as the best model for each defensin. Because of the low sequence identity (<30%) between EfAMP-1 and proteins deposited in the PDB, *LOMETS* (LOcal Meta-Threading-Server) [64] was used to determine the best template/alignment, which applies a threading/fold recognition approach. Following, 3D models of EfAMP-1 were generated using *Modeller* v9.19 [63] and processed as described above. LsAMP-1, LsAMP-2, and EfAMP-1 best models were quality checked using *ProSa* II [65] and *MolProbity [66]* and visualized using *PyMOL* v2.1 (https://pymol.org/2/).

### - Phylogenetic analyses

*Dali* server [67] was used to perform pairwise structural alignments of LsAMP-1, LsAMP-2, and EfAMP-1 with protein structures available in PDB; and neighbor-joining trees were subsequently inferred using these alignments with *SeaView* v4.6 [68].

### - Molecular dynamics

Molecular dynamics simulations with LsAMP-1, LsAMP-2, and EfAMP-1 were performed using *Gromacs* v5 [69] applying the all-atom force field OPSL [70]. First of all, models were minimized using the steepest *descent* algorithm, centered in dodecahedral boxes with sides of 2.0 nm filled with water molecules that were modeled with TIP3P (TIP 3-point) and their geometry constrained with SETTLE algorithm [71]. Then, systems were neutralized with 0.15 mol/L of sodium chloride and after applying position constraints, equilibrated using NVT followed by NPT ensembles, considering a temperature of 300 K and pressure of 1 bar, both for 100 ps. All-atom bond lengths were linked using the *LINCS* algorithm [72]; electrostatic interactions were treated with the particle-mesh Ewald (PME) method [73], considering the electrostatic and van der Waals cutoffs of 1.0 nm, updating the list of neighbors of each atom every 10 simulation steps of 2 fs. Finally, models were released from their position constraints and production was performed for 1 μs using the *leap-frog* algorithm as the integrator. These MD simulations were analyzed by means of root-mean-square deviation (RMSD), using the backbones as units for root square calculations, and by means of Dictionary of Protein Secondary Structures (DSSP), using rms and do_dssp available from *Gromacs* package, respectively.

### - Ligand prediction

To shed some light into the mechanism of action of these defensins, comparative structural analysis were performed, using *Cofactor [74]*, in which data were further confirmed through molecular docking and MD simulations. Predicted potential ligands suggested during *Cofactor* analysis were docked to the predicted binding site within the defensins, using *DockThor* web-server (https://www.dockthor.lncc.br/v2/). The 2D representations of all tested ligands were retrieved from *PubChem* (https://pubchem.ncbi.nlm.nih.gov/) and *obabel* [75] was used to convert them to 3D structures. Next, protonation states of the three defensins were automatically predicted using H++ (http://biophysics.cs.vt.edu/H++), applying pH = 7.5, octahedral solvent box, TIP3P water model, and neutralization with NaCl. Finally, Charmm force field parameters of potential cellular ligands, were generated with *SwissParam* (https://www.swissparam.ch/), then, predicted defensin-ligand pairs were subjected to 200 ns MD simulations using all-atom force field Charmm27 [76] and following the procedure described in the previous section. Binding pose 2D representations were done using Pose2view (https://proteins.plus/).

## Ethics approval and consent to participate

Not applicable

## Consent for publication

Not applicable

## Availability of data and materials

The datasets used and/or analysed during the current study are available from the corresponding author on reasonable request.

## Competing interests

The author has no competing interests to declare that are relevant to the content of this article.

## Authors’ contributions

Not applicable

## Acknowledgements

This work was financially by “Fundação de Amparo à Pesquisa do Estado de São Paulo” (FAPESP) (process 0022/00538-0).

---

**Suplemental Material 1.**
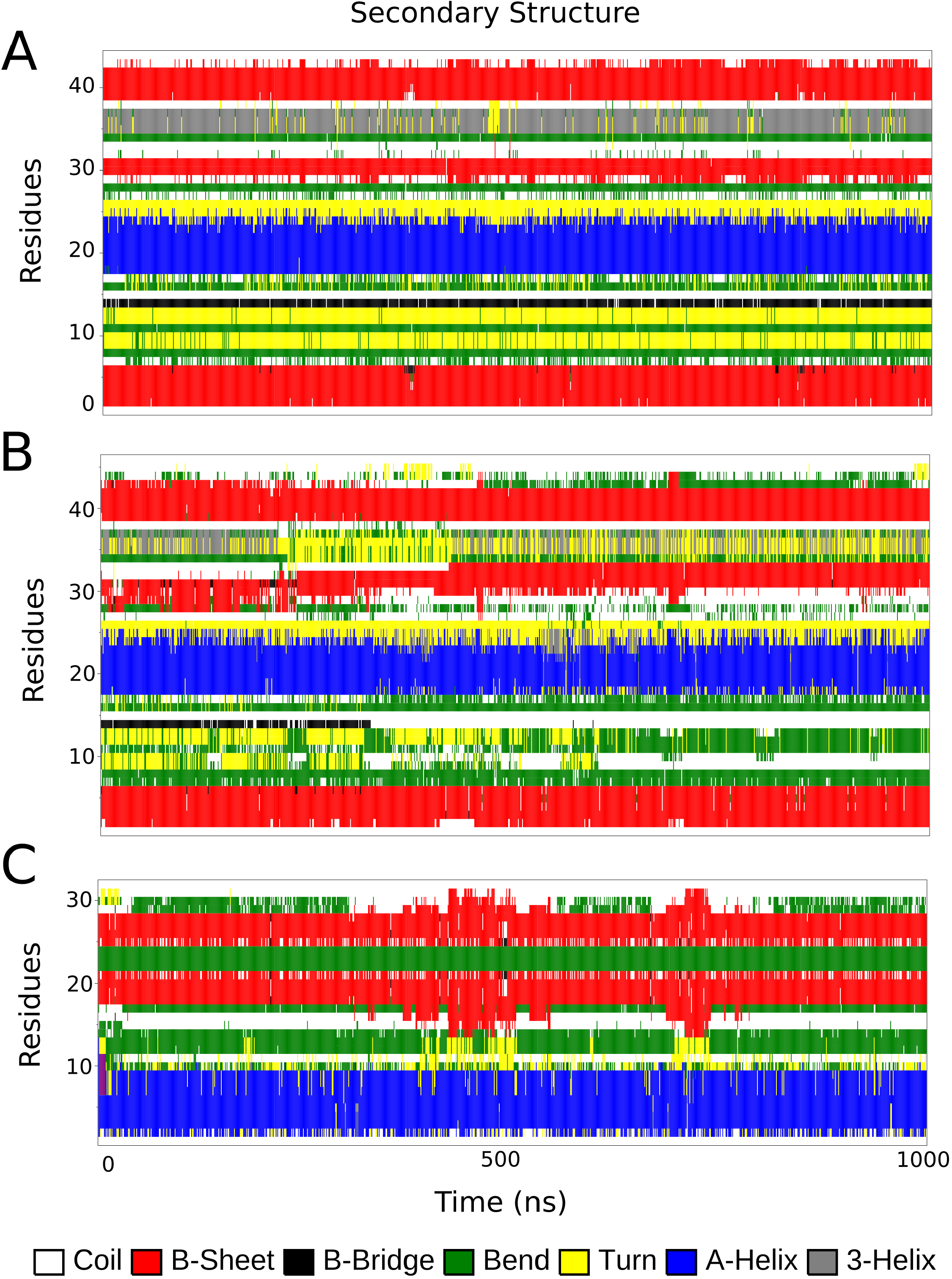
Secondary structure analyses (DSSP) of the models (A) LsAMP1 (B) LsAMP2 (C) EfAMPl over the molecular dynamics run of 1 microsecond.

## Notes

### Competing Interest Statement

The authors have declared no competing interest.

## References

1. Davies J, Davies D. Origins and Evolution of Antibiotic Resistance. Microbiol Mol Biol Rev. 2010;74:417–33.

2. Levy SB, Bonnie M. Antibacterial resistance worldwide: Causes, challenges and responses. Nat Med. 2004;10:S122–9.

3. Barber M, Rozwadowska-Dowzenko M. Infection By Penicillin-Resistant Staphylococci. Lancet. 1948;252:641–4.

4. Small PM, Shafer RW, Hopewell PC, Singh SP, Murphy MJ, Desmond E, et al. Exogenous reinfection with multidrug-resistant Mycobacterium tuberculosis in patients with advanced HIV infection. N Engl J Med. 1993;328:1137–44.

5. Nguyen S V., McShan WM. Chromosomal islands of Streptococcus pyogenes and related streptococci: molecular switches for survival and virulence. Front Cell Infect Microbiol. 2014;4 August:1–14.

6. Muto CA, Jernigan JA, Ostrowsky BE, Richet HM, Jarvis WR, Boyce JM, et al. SHEA guideline for preventing nosocomial transmission of multidrug‐resistant strains of Staphylococcus aureus and Enterococcus. Infect Control Hosp Epidemiol. 2003;24:362–86.

7. Fernández A, Pereira MJ, Suárez JM, Poza M, Treviño M, Villalón P, et al. Emergence in Spain of a multidrug-resistant Enterobacter cloacae clinical isolate producing SFO-1 extended-spectrum β-lactamase. J Clin Microbiol. 2011;49:822–8.

8. Aloush V, Navon-venezia S, Seigman-igra Y, Cabili S, Carmeli Y. Multidrug-Resistant Pseudomonas aeruginosa : Risk Factors and Clinical Impact. Society. 2006;50:43–8.

9. Gulshan K, Moye-Rowley WS. Multidrug resistance in fungi. Eukaryot Cell. 2007;6:1933–42.

10. Bontems F, Roumestand C, Gilquin B, Menez A, Toma F. Refined structure of charybdotoxin: common motifs in scorpion toxins and insect defensins. Science (80-). 1991;254:1521–3.

11. Mygind PH, Fischer RL, Schnorr KM, Hansen MT, Sönksen CP, Ludvigsen S, et al. Plectasin is a peptide antibiotic with therapeutic potential from a saprophytic fungus. Nature. 2005;437:975–80.

12. Lacerda AF, Vasconcelos ÉAR, Pelegrini PB, Grossi de Sa MF. Antifungal defensins and their role in plant defense. Front Microbiol. 2014;5 APR:1–10.

13. Da Silva P, Jouvensal L, Lamberty M, Bulet P, Caille A, Vovelle F. Solution structure of termicin, an antimicrobial peptide from the termite Pseudacanthotermes spiniger. Protein Sci. 2003;12:438–46.

14. Roch P, Yang Y, Toubiana M, Aumelas A. NMR structure of mussel mytilin, and antiviral-antibacterial activities of derived synthetic peptides. Dev Comp Immunol. 2008;32:227–38.

15. Johnson C, Kleshchenko Y, Furtak V, Nde P, Pratap S, Madison N, et al. Trypanosoma cruzi regulates the defensin alpha-1 interactome network and defensin alpha-1 causes trypanosome membrane pore formation to control the early process of infection. J Immunol. 2011;186 1 Supplement:56.30 LP–56.30.

16. Spelbrink RG. Differential Antifungal and Calcium Channel-Blocking Activity among Structurally Related Plant Defensins. Plant Physiol. 2004;135:2055–67.

17. Yang WS, Ko J, Kim E, Kim JH, Park JG, Sung NY, et al. 21-O-angeloyltheasapogenol E3, a novel triterpenoid saponin from the seeds of tea plants, inhibits macrophage-mediated inflammatory responses in a NF-B-dependent manner. Mediators Inflamm. 2014;2014.

18. Liu Y-J, Cheng C-S, Lai S-M, Hsu M-P, Chen C-S, Lyu P-C. Solution structure of the plant defensin VrD1 from mung bean and its possible role in insecticidal activity against bruchids. Proteins Struct Funct Bioinforma. 2006;63:777–86.

19. Wijaya R, Neumann GM, Condron R, Hughes AB, Polya GM. Defense proteins from seed of Cassia fistula include a lipid transfer protein homologue and a protease inhibitory plant defensin. Plant Sci. 2000;159:243–55.

20. Kudryashova E, Quintyn R, Seveau S, Lu W, Wysocki VH, Kudryashov DS. Human defensins facilitate local unfolding of thermodynamically unstable regions of bacterial protein toxins. Immunity. 2014;41:709–21.

21. Dias RDO, Franco OL. Cysteine-stabilized αβ defensins: From a common fold to antibacterial activity. Peptides. 2015;72:64–72.

22. Shahabuddin M, Fields I, Bulet P, Hoffmann JA, Miller LH. Plasmodium gallinaceum: Differential killing of some mosquito stages of the parasite by insect defensin. Exp Parasitol. 1998;89:103–12.

23. Boulanger N, Boulanger N, Lowenberger C, Lowenberger C, Volf P, Volf P, et al. Characterization of a Defensin from the Sand Fly. Microbiology. 2004;72:7140–6.

24. Lamberty M, Caille A, Landon C, Tassin-Moindrot S, Hetru C, Bulet P, et al. Solution structures of the antifungal heliomicin and a selected variant with both antibacterial and antifungal activities. Biochemistry. 2001;40:11995–2003.

25. Vizioli J, Richman AM, Uttenweiler-Joseph S, Blass C, Bulet P. The defensin peptide of the malaria vector mosquito anopheles gambiae: Antimicrobial activities and expression in adult mosquitoes. Insect Biochem Mol Biol. 2001;31:241–8.

26. Lynn D. The Ciliated Protozoa. 3rd edition. New York, NY: Pergamon Press; 2008.

27. Zhu S, Gao B, Tytgat J. Phylogenetic distribution, functional epitopes and evolution of the CSαβ superfamily. Cell Mol Life Sci. 2005;62:2257–69.

28. Landon C. Lead optimization of antifungal peptides with 3D NMR structures analysis. Protein Sci. 2004;13:703–13.

29. Yount NY, Yeaman MR. Multidimensional signatures in antimicrobial peptides. Proc Natl Acad Sci. 2004;101:7363–8.

30. Yount NY, Andrés MT, Fierro JF, Yeaman MR. The γ-core motif correlates with antimicrobial activity in cysteine-containing kaliocin-1 originating from transferrins. Biochim Biophys Acta - Biomembr. 2007;1768:2862–72.

31. Fant F, Vranken W, Broekaert W, Borremans F. Determination of the three-dimensional solution structure of Raphanus sativus antifungal protein 1 by 1H NMR. J Mol Biol. 1998;279:257–70.

32. Zhu S. Evidence for myxobacterial origin of eukaryotic defensins. Immunogenetics. 2007;59:949–54.

33. Zhu S. Discovery of six families of fungal defensin-like peptides provides insights into origin and evolution of the CSαβ defensins. Mol Immunol. 2008;45:828–38.

34. Koehbach J. Structure-Activity Relationships of Insect Defensins. Front Chem. 2017;5 July:1–10.

35. Valachova I, Prochazka E, Bohova J, Novak P, Takac P, Majtan J. Antibacterial properties of lucifensin in Lucilia sericata maggots after septic injury. Asian Pac J Trop Biomed. 2014;4:358–61.

36. Wu J, Liu S, Wang H. Invasive fungi-derived defensins kill drug-resistant bacterial pathogens. Peptides. 2018;99 November 2017:82–91.

37. Valore E V, Ganz T. Laboratory production of antimicrobial peptides in native conformation. Antibact Pept Protoc. 1997;78:115–31.

38. Savoia D, Avanzini C, Allice T, Callone E, Guella G, Dini F. Antimicrobial Activity of Euplotin C, the Sesquiterpene Taxonomic Marker from the Marine Ciliate Euplotes crassus. Antimicrob Agents Chemother. 2004;48:3828–33.

39. Lobban CS, Hallam SJ, Mukherjee P, Petrich JW. Photophysics and multifunctionality of hypericin-like pigments in heterotrich ciliates: A phylogenetic perspective. Photochem Photobiol. 2007;83:1074–94.

40. Petrelli D, Buonanno F, Vitali LA, Ortenzi C. Antimicrobial activity of the protozoan toxin climacostol and its derivatives. Biologia (Bratisl). 2012;67:525–9.

41. Cui P, Dong Y, Li Z, Zhang Y, Zhang S. Identification and functional characterization of an uncharacterized antimicrobial peptide from a ciliate Paramecium caudatum. Dev Comp Immunol. 2016;60:53–65.

42. Brogden KA. Antimicrobial peptides: Pore formers or metabolic inhibitors in bacteria? Nat Rev Microbiol. 2005;3:238–50.

43. Xu XX, Zhang YQ, Freed S, Yu J, Gao YF, Wang S, et al. An anionic defensin from Plutella xylostella with potential activity against Bacillus thuringiensis. Bull Entomol Res. 2016;106:790–800.

44. Wen H, Lan X, Cheng T, He N, Shiomi K, Kajiura Z, et al. Sequence structure and expression pattern of a novel anionic defensin-like gene from silkworm (Bombyx mori). Mol Biol Rep. 2009;36:711–6.

45. Lai R, Liu H, Hui Lee W, Zhang Y. An anionic antimicrobial peptide from toad Bombina maxima. Biochem Biophys Res Commun. 2002;295:796–9.

46. Becucci L, Valensin D, Innocenti M, Guidelli R. Dermcidin, an anionic antimicrobial peptide: Influence of lipid charge, pH and Zn2+ on its interaction with a biomimetic membrane. Soft Matter. 2014;10:616–26.

47. Contreras G, Santhosh Braun M, Schä H, Wink M. Recombinant AfusinC, an anionic fungal CSαβ defensin from Aspergillus fumigatus, exhibits antimicrobial activity against gram-positive bacteria. 2018. https://doi.org/10.1371/journal.pone.0205509.

48. Katan M, Cockcroft S. Phosphatidylinositol(4,5)bisphosphate: diverse functions at the plasma membrane. Essays Biochem. 2020;64:513.

49. Yoneda A, Kanemaru K, Matsubara A, Takai E, Shimozawa M, Satow R, et al. Phosphatidylinositol 4,5-bisphosphate is localized in the plasma membrane outer leaflet and regulates cell adhesion and motility. 2020;527:1050–6.

50. Poon IKH, Baxter AA, Lay FT, Mills GD, Adda CG, Payne JAE, et al. Phosphoinositide-mediated oligomerization of a defensin induces cell lysis. Elife. 2014;2014.

51. Benson DA, Cavanaugh M, Clark K, Karsch-Mizrachi I, Lipman DJ, Ostell J, et al. GenBank. Nucleic Acids Res. 2012;41:D36–42.

52. Rice P, Longden I, Bleasby A. EMBOSS: the European Molecular Biology Open Software Suite. Trends Genet. 2000;16:276–7.

53. Dutilh BE, Jurgelenaite R, Szklarczyk R, van Hijum SAFTFT, Harhangi HR, Schmid M, et al. FACIL: Fast and accurate genetic code inference and logo. Bioinformatics. 2011;27:1929–33.

54. Camacho C, Coulouris G, Avagyan V, Ma N, Papadopoulos J, Bealer K, et al. BLAST+: architecture and applications. BMC Bioinformatics. 2009;10:421.

55. Boutet E, Lieberherr D, Tognolli M, Schneider M, Bansal P, Bridge AJ, et al. UniProtKB/Swiss-Prot, the Manually Annotated Section of the UniProt KnowledgeBase: How to Use the Entry View. Humana Press, New York, NY; 2016. p. 23–54.

56. Almagro Armenteros JJ, Tsirigos KD, Sønderby CK, Petersen TN, Winther O, Brunak S, et al. SignalP 5.0 improves signal peptide predictions using deep neural networks. Nat Biotechnol. 2019;37:420–3.

57. Käll L, Krogh A, Sonnhammer EL. L. A combined transmembrane topology and signal peptide prediction method. J Mol Biol. 2004;338:1027–36.

58. McGuffin LJ, Bryson K, Jones DT. The PSIPRED protein structure prediction server. Bioinformatics. 2000;16:404–5.

59. Lee HT, Lee CC, Yang JR, Lai JZC, Chang KY, Ray O. A large-scale structural classification of Antimicrobial peptides. Biomed Res Int. 2015;2015.

60. Waghu FH, Barai RS, Gurung P, Idicula-Thomas S. CAMPR3: A database on sequences, structures and signatures of antimicrobial peptides. Nucleic Acids Res. 2016;44:D1094–7.

61. Xiao X, Wang P, Lin WZ, Jia JH, Chou KC. IAMP-2L: A two-level multi-label classifier for identifying antimicrobial peptides and their functional types. Anal Biochem. 2013;436:168–77.

62. Zimmermann L, Stephens A, Nam S-ZZ, Rau D, Kübler J, Lozajic M, et al. A Completely Reimplemented MPI Bioinformatics Toolkit with a New HHpred Server at its Core. J Mol Biol. 2017;430:2237–43.

63. Webb B, Sali A. Comparative protein structure modeling using MODELLER. Curr Protoc Bioinforma. 2016;2016 June:5.6.1–5.6.37.

64. Wu S, Zhang Y. LOMETS: A local meta-threading-server for protein structure prediction. Nucleic Acids Res. 2007;35:3375–82.

65. Sippl MJ. Recognition of errors in three-dimensional structures of proteins. Proteins Struct Funct Genet. 1993;17:355–62.

66. Chen VBVBVB, Arendall WBB, Headd JJJ, Keedy DADADA, Immormino RMRM, Kapral GJGJ, et al. MolProbity: All-atom structure validation for macromolecular crystallography. Acta Crystallogr Sect D Biol Crystallogr. 2010;66:12–21.

67. Holm L, Laakso LM. Dali server update. Nucleic Acids Res. 2016;44:W351–5.

68. Gouy M, Guindon S, Gascuel O. SeaView Version 4: A Multiplatform Graphical User Interface for Sequence Alignment and Phylogenetic Tree Building. Mol Biol Evol. 2010;27:221–4.

69. Abraham MJ, Murtola T, Schulz R, Páll S, Smith JC, Hess B, et al. Gromacs: High performance molecular simulations through multi-level parallelism from laptops to supercomputers. SoftwareX. 2015;1–2:19–25.

70. Jorgensen WL, Maxwell DS, Tirado-Rives J. Development and testing of the OPLS all-atom force field on conformational energetics and properties of organic liquids. J Am Chem Soc. 1996;118:11225–36.

71. Miyamoto S, Kollman PA. Settle: An analytical version of the SHAKE and RATTLE algorithm for rigid water models. J Comput Chem. 1992;13:952–62.

72. Hess B, Bekker H, Berendsen HJC, Fraaije JGEM. LINCS: a linear constraint solver for molecular simulations. J Comput Chem. 1997;18:1463–72.

73. Darden T, York D, Pedersen L. Particle mesh Ewald: An N·log(N) method for Ewald sums in large systems. J Chem Phys. 1993;98:10089–92.

74. Zhang C, Freddolino PL, Zhang Y. COFACTOR: Improved protein function prediction by combining structure, sequence and protein-protein interaction information. Nucleic Acids Res. 2017;45:W291–9.

75. O’Boyle NM, Banck M, James CA, Morley C, Vandermeersch T, Hutchison GR. Open Babel: An open chemical toolbox. J Cheminform. 2011;3:33.

76. Foloppe N, MacKerell, Jr. AD. All-atom empirical force field for nucleic acids: I. Parameter optimization based on small molecule and condensed phase macromolecular target data. J Comput Chem. 2000;21:86–104.

